# Increased autophagy in ephrinB2 deficient osteocytes is associated with hypermineralized, brittle bones

**DOI:** 10.1101/260711

**Authors:** Christina Vrahnas, Toby A Dite, Niloufar Ansari, Blessing Crimeen-Irwin, Huynh Nguyen, Mark R Forwood, Yifang Hu, Mika Ikegame, Keith R Bambery, Cyril Petibois, Mark J Tobin, Gordon K Smyth, Jonathan S Oakhill, T John Martin, Natalie A Sims

## Abstract

Mineralized bone forms when collagen-containing osteoid accrues hydroxyapatite crystals. This process has two phases: a rapid initiation (primary mineralization), followed by slower accrual of mineral (secondary mineralization) that continues until that portion of bone is renewed by remodelling. Within the bone matrix is an interconnected network of cells termed osteocytes. These cells are derived from bone-forming osteoblasts. Osteoblast differentiation requires expression of ephrinB2, and we were intrigued about why ephrinB2 continues to be expressed in mature osteocytes. To determine its function in osteocytes, we developed an osteocyte-specific ephrinB2 null mouse and found they exhibited a brittle bone phenotype. This was not caused by a change in bone mass, but by an intrinsic defect in the strength of the bone material. Although the initiation of osteoid mineralization occurred at a normal rate, the process of secondary mineralization was accelerated in these mice. The maturing mineralized bone matrix incorporated mineral and carbonate more rapidly than controls, indicating that osteocytic ephrinB2 suppresses mineral accumulation in bone. No known regulators of mineralization were modified in the bone of these mice. However, RNA sequencing showed differential expression of a group of autophagy-associated genes, and increased autophagic flux was confirmed in ephrinB2 knockdown osteocytes. This indicates that the process of secondary mineralization in bone makes use of autophagic machinery in a manner that is limited by ephrinB2 in osteocytes, and that this process may be disrupted in conditions of bone fragility.

## Introduction

The skeleton is unique in the human body because its organic structure is hardened by integration of mineral, allowing it to provide support for locomotion and protection for internal organs. This mineralized bone forms when collagen-containing osteoid, deposited by osteoblasts, accrues hydroxyapatite crystals. The process has two phases: a rapid initiation (primary mineralization), followed by slower accrual of mineral (secondary mineralization) that continues until that portion of bone is renewed by remodelling. During bone formation, osteoblasts become incorporated in the non-mineralized osteoid and differentiate into osteocytes to form a highly connected network of specialized cells residing within the mineralized matrix^1,2^. In response to stimuli such as mechanical load, hormones, and cytokines, osteocytes release proteins that both stimulate^3^ and inhibit^4^ bone forming osteoblasts. Both osteoblasts and osteocytes express proteins that initiate mineralization of the osteoid matrix ^4‐7^, but it is not known how the speed of continuing mineral deposition is controlled.

Two related agents that stimulate bone formation are the agonists of the parathyroid hormone (PTH) receptor (PTHR1): PTH, and PTH-related protein (PTHrP). Locally-derived PTHrP, from osteoblasts and osteocytes stimulates bone formation in a paracrine manner ^8,9^, and this action has been exploited by the pharmacological agents teriparatide and abaloparatide. These are the only pharmacological agents that are currently available clinically that can increase bone mass in patients with fragility fractures ^10,11^. Both PTH and PTHrP treatment substantially increase the expression of the contact-dependent signalling molecule ephrinB2 in osteoblasts *in vitro* and in bone *in vivo* ^12^. We have previously shown that the interaction of ephrinB2 (gene name: *Efnb2*) with its receptor, EphB4, provides a checkpoint through which osteoblasts must pass to reach late stages of osteoblast and osteocyte differentiation ^13‐16^. Furthermore, osteoblast-specific deletion of ephrinB2 compromised bone strength by impairing osteoblast differentiation, and delaying the initiation of bone mineralization, ultimately leading to a mild osteomalacia (high osteoid content)^14^. This indicated that osteoid mineralization is initiated by osteoblasts after the ephrinB2:EphB4 differentiation checkpoint.

We were intrigued that ephrinB2 expression remains high in fully embedded osteocytes ^12^, beyond the differentiation checkpoint. Given the extensively connected nature of osteocytes and the contact-dependent nature of ephrinB2:EphB4 signalling, we hypothesized that ephrinB2 regulates the function of osteocytes embedded in the bone matrix. We undertook the present work to determine the requirement for ephrinB2 expression in osteocytes. We find that ephrinB2 in osteocytes does not regulate the initiation of mineralization, but is required to limit secondary mineral accrual and retain the flexibility of the bone matrix. We also show that, in the absence of ephrinB2, osteocytes exhibit modified levels of genes associated with autophagy and increased autophagic flux, providing evidence that autophagic processes in osteocytes directly control mineralization.

## Results

### EphrinB2 is upregulated by PTH and PTHrP in 0cy454 cells

Upregulation *Efnb2* in osteocytes by PTH and PTHrP stimulation was confirmed in 0cy454 cells (Fig. 1A); PTH(1-34) and PTHrP(1-141) both significantly increased *Efnb2* mRNA levels. shRNA knockdown of PTHrP (*Pthlh*) in 0cy454 cells resulted in significantly lower *Efnb2* mRNA levels at all timepoints, compared to vector control (Fig. 1B), indicating that endogenous PTHrP expression by osteocytes is required for normal *Efnb2* expression in osteocytes.

**Figure 1:**
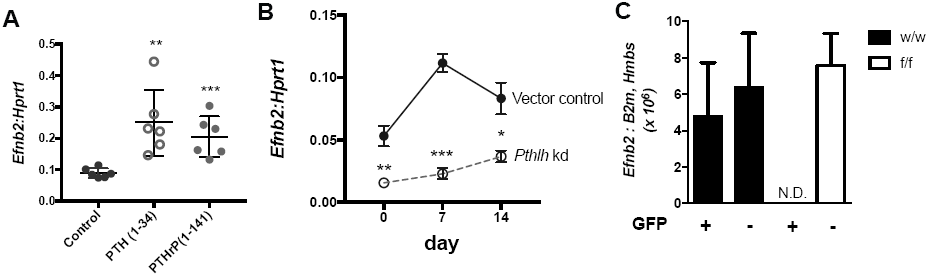
*Efnb2* is upregulated by PTH and PTHrP in osteocyte-like cells and confirmation of ephrinB2 knockdown in *Dmp1Cre.EfnB2*^*f/f*^ osteocytes. **A,B:** *Efnb2* mRNA levels, relative to *Hprtl,* in 0cy454 cells differentiated for 14 days and treated with lOnM hPTH(l-34) or hPTHrP(l-141) for 6h **(A)** and 0cy454 cells with stable shRNA knockdown of PTHrP (*Pthlh*) at 0, 7 and 14 days of differentiation **(B).** Data are mean ± SEM, n=6 replicates; representative of 3 independent experiments. **C:** *Efnb2* mRNA levels in GFP+ osteocytes isolated from *Dmp1Cre.DMPl-GFP-Tg.Efnb2*^*w*^*/*^*w*^ (w/w) and *Dmp1Cre.DMPl-GFP-Tg.Efnb2*^*f/f*^ [f/f) mice compared to GFP-cells. N.D. = not detected. Data mean ± SEM of 2 experiments, n= 6 mice/group; pooled. *\*p<0.05, **p<0.01, ***p<0.001* compared to control by Student’s t-test.

### Deletion of ephrinB2 in osteocytes results in a brittle bone phenotype

FACS purification of DMP1-GFP+ osteocytes and use of primers targeted to the exon 1-2 boundary of *Efnb2* confirmed that *Efnb2* mRNA was present in osteocytes of control mice. Analysis of GFP+ cells confirmed effective targeting of *Efnb2* in osteocytes from *Dmp1Cre.Efnb2*^*f/f*^ mice, while retaining *Efnb2* in non-osteocytic cells (Fig. 1C).

Three-point bending tests demonstrated a strength defect in female *Dmp1Cre.Efnb2*^*f/f*^ femora compared to *Dmp1Cre.Efnb2*^*w*^*/*^*w*^ controls (Fig. 2A-F, Table 1). There was no significant difference in bone strength observed in males (Supplementary Table 1). A significantly higher percentage of bones from *Dmp1Cre.Efnb2*^*f/f*^ mice fractured at lower deformation than controls (Fig. 2B). Average load-deformation curves also showed this (Fig. 2A), and indicated that femora from *Dmp1Cre.Efnb2*^*f/f*^ mice could withstand less force before yielding (Table 1) and before breaking (ultimate force, Fig. 2C), and deformed less at the maximum force that bones could withstand, compared to controls (Fig. 2D). The reduction in ultimate deformation was largely due to a deficit in the extent of deformation from the yield point onwards (Fig. 2E, Table 1). Energy absorbed to failure was also significantly lower in *Dmp1Cre.Efnb2*^*f/f*^ mice compared to controls (Fig. 2F). Stiffness (the slope of the load / deformation curve before the yield point) was not significantly modified (Table 1). Instead, *Dmp1Cre.Efnb2*^*f/f*^ femora had a lower yield point, and once they began to yield, they withstood less force and deformed less than controls.

**Figure 2:**
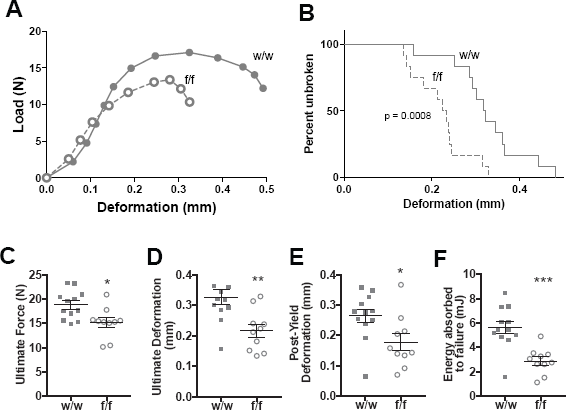
12-week-old female *Dmp1Cre.EfnB2*^*f/f*^ (f/f) mice have impaired femoral strength compared to *Dmp1Cre* controls (w/w). A: Average Load-Deformation curve from 3 point bending tests (each dot represents the average load and deformation for the noted sample group; error bars excluded to highlight the shape of curves). B: Kaplan-Meier curve showing percentage of unbroken femora with increasing deformation (p<0.0008 based on Log-rank (Mantel-Cox) test). C-F: Calculated indices of bone strength: ultimate force (C), ultimate deformation (D), post-yield deformation (E) and energy absorbed to failure (F). Data are mean ± SEM, n = 10-12/group. *p<0.05, **p<0.01, ***p<0.001 vs w/w controls.

**Table 1.** Additional structural and material properties of 12-week old female *Dmp1Cre.Efnb2*^*f/f*^ *femoral* cortical bone, compared to controls.

|  | *Dmp1Cre.Efnb2<sup>w/w</sup>* | *Dmp1Cre.Efnb2<sup>f/f</sup>* |
| --- | --- | --- |
| **Structural properties** |  |  |
| Yield Force (N) | 14.98 ± 0.63 | 11.15 ± 0.90<sup>b</sup> |
| Yield deformation (mm) | 0.18 ± 0.01 | 0.12 ± 0.01<sup>b</sup> |
| Stiffness (N/mm) | 139.32 ± 10.16 | 134.59 ± 10.31 |
| Anteroposterior (AP) width (mm) | 1.24 ± 0.02 | 1.25 ± 0.02 |
| Mediolateral (ML) width (mm) | 1.80 ± 0.03 | 1.75 ± 0.01 |
| **Material properties** |  |  |
| Yield Stress (MPa) | 31.96 ± 2.12 | 24.15 ± 1.81<sup>a</sup> |
| Yield Strain (%) | 0.04 ± 0.002 | 0.02 ± 0.001<sup>b</sup> |
| Elastic Modulus (MPa) | 1430.5 ± 105.8 | 1320.4 ± 93.5 |
| **Additional data** |  |  |
| Periosteal Perimeter (µm) | 6.90 ± 0.13 | 6.86 ± 0.07 |
| Marrow Area (mm<sup>2</sup>) | 0.94 ± 0.04 | 0.97 ± 0.02 |
| Periosteal Mineral Appositional Rate (µm/day) | 0.98 ± 0.12 | 1.07 ± 0.10 |
| Periosteal Mineralising Surface / Bone Surface (%) | 88.41 ± 7.45 | 80.82 ± 6.30 |
| Cortical Tissue Mineral Density (g/cm<sup>3</sup>) | 1.13 ± 0.02 | 1.13 ± 0.02 |

Data are represented as mean ± SEM; n = 12/group
<sup>a</sup>p < 0.05, <sup>b</sup>p < 0.01, versus *Dmp1Cre.Efnb2<sup>w/w</sup>* controls, Student's t-test.

The reduction in bone strength in *Dmp1Cre.Efnb2*^*f/f*^ mice was not associated with any significant difference in moment of inertia (Fig. 3A) or dimensions of the cortical bone (Table 1). Indeed, when normalised for cortical dimensions, femora from *Dmp1Cre.Efnb2*^*f/f*^ mice showed impaired material strength compared to controls (Fig. 3B), including lower yield stress and strain (Table 1), lower ultimate stress (Fig. 3C), lower ultimate strain (Fig. 3D), and lower toughness (Fig. 3E). Elastic modulus was not significantly altered (Table 1), consistent with the normal stiffness data obtained. Reference point indentation also indicated a change in material strength independent of geometry, with a significantly greater indentation distance increase (IDI) (the increase in the test probe’s indentation distance in the last cycle relative to the first cycle) in *Dmp1Cre.Efnb2*^*f/f*^ bones compared to controls (Fig. 3F). No other parameters measured by RPI were significantly altered (Supplementary Table 2).

**Figure 3:**
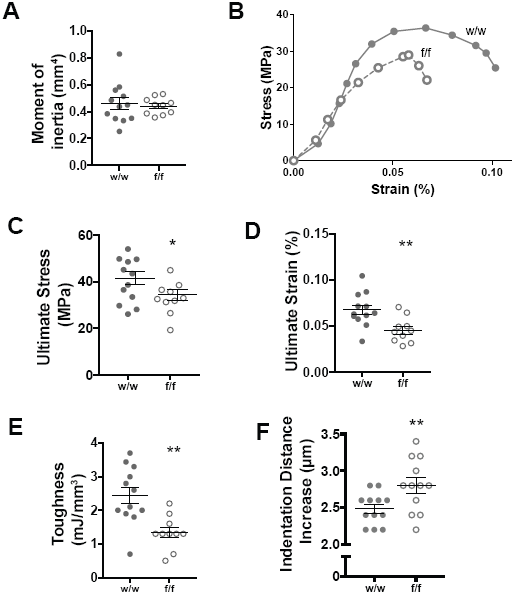
Impaired strength in *Dmp1Cre.Efnb2*^f/f^ mice is a material defect. **A:** Femoral moment of inertia. **B-E:**3 point bending test data corrected for bone cross sectional area, including average stress-strain curve (each dot represents the average load and deformation for the noted sample group; error bars excluded to highlight the shape of curves) (B), ultimate stress (C), ultimate strain (D), and toughness (E) of 12-week old female *Dmp1Cre.Efnb2*^*w*^*/*^*w*^ (w/w) and *Dmp1Cre.Efnb2*^*f/f*^ (f/f) femora. **F:** Indentation distance increase (IDI) derived from reference point indentation of femora. Data are represented as mean ± SEM, n = 10-12/group. *p<0.05, **p<0.01 vs w/w controls.

**Table 2.** Histomorphometric analyses of trabecular bone in 12-week old female *Dmp1Cre.Efnb2*^*f/f*^ tibiae compared to sex and age-matched *Dmp1Cre.Efnb2*^*w*^*/*^*w*^ controls.

|  | *Dmp1Cre.Efnb2<sup>w/w</sup>* | *Dmp1Cre.Efnb2<sup>f/f</sup>* |
| --- | --- | --- |
| Mineral Apposition Rate ($\mu\text{m}/\text{day}$) | 2.09 ± 0.09 | 1.96 ± 0.10 |
| Mineralising surface/Bone Surface (%) | 34.51 ± 2.52 | 34.87 ± 1.04 |
| Bone Formation Rate/Bone Surface ($\mu\text{m}^2/\mu\text{m}/\text{day}$) | 0.71 ± 0.04 | 0.68 ± 0.04 |
| Osteoid Thickness ($\mu\text{m}$) | 2.26 ± 0.17 | 2.66 ± 0.21 |
| Osteoid Surface (%) | 7.39 ± 1.45 | 9.83 ± 2.29 |
| Osteoblast Surface (%) | 13.60 ± 1.81 | 16.25 ± 2.20 |
| Osteoclast Surface (%) | 8.78 ± 1.25 | 11.58 ± 0.95 |

Data are represented as mean ± SEM; n = 7-11/group

This indicates that *Dmp1Cre.Efnb2*^*f/f*^ bones are more brittle than controls due to a defect in bone material compisition.

### Bone formation rate is unchanged, but osteocyte density is greater in Dmp1Cre.Efnb2^f/f^ bones

In contrast to *OsxCre.Efnb2*^*f/f*^ mice (with ephrinB2 deleted in osteoblasts) which exhibited reduced mineral appositional rate at the periosteum ^*14*^, *Dmp1Cre.Efnb2*^*f/f*^ mice exhibited no significant change in periosteal mineral appositional rate (Table 1) or periosteal mineralising surface (Table 1). We also assessed trabecular bone, which was of normal volume (Supplementary Table 3), and detected no significant difference in trabecular bone formation rate, mineral appositional rate, or mineralising surface (Table 2). There were also no differences detected in osteoid thickness, osteoid surface, osteoblast surface or osteoclast surface in *Dmp1Cre.Efnb2*^*f/f*^ bones compared to controls (Table 2). This indicated that osteoblast-mediated osteoid deposition and the initiation of bone mineralization occurred at a normal rate in *Dmp1Cre.Efnb2*^*f/f*^ bones.

**Table 3.**
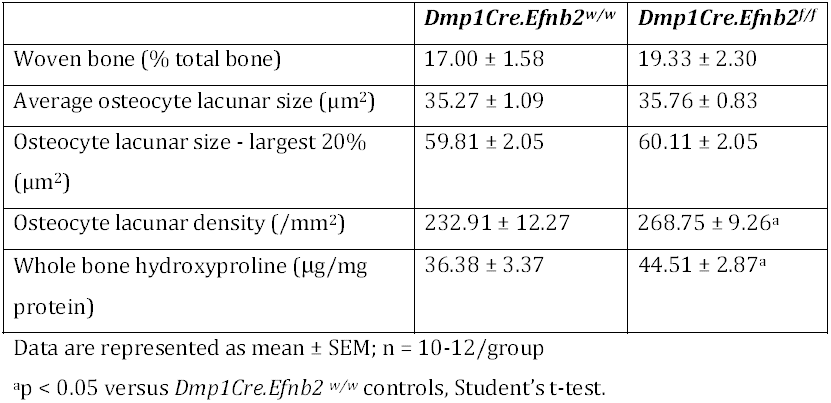
Collagen fibre deposition and osteocyte lacunar parameters in the femoral midshaft from 12-week old female *Dmp1Cre.Efnb2*^*w*^*/*^*w*^ (w/w) and *Dmp1Cre.Efnb2*^*f/f*^ (f/f) mice.

Backscattered electron microscopy detected no change in osteocyte lacunar size in the femoral midshaft of *Dmp1Cre.Efnb2*^*f/f*^ mice, however osteocyte lacunar density was significantly greater compared to controls (Table 3). This suggests a greater rate of osteocyte incorporation into the bone matrix during osteoid deposition.

### Dmp1Cre.Efnb2f/f bone has greater mineral and carbonate deposition

Since material strength was impaired and the (blinded) technician cutting the sections for histomorphometry noted that some samples were very difficult to cut, we assessed bone material by a number of methods. We detected no difference in the proportion of woven to lamellar cortical bone by polarised light microscopy (Table 3) and no significant alteration in cortical tissue mineral density (Table 1) in *Dmp1Cre.Efnb2*^*f/f*^ femora compared to controls. We then used sFTIRM to measure bone composition at the periosteum, a region lacking bone remodelling, thereby allowing assessment of the bone matrix as mineralization progresses ^17^ Three regions of increasing matrix maturity were measured (Fig. 4A). Average spectra for each genotype, taken from the intermediate region indicated altered spectral geometry in *Dmp1Cre.Efnb2*^*f/f*^ bone compared to controls (Fig. 4B). *Dmp1Cre.Efnb2*^*f/f*^ bone had higher phosphate and carbonate peaks indicating a greater level of mineralization. In addition, *Dmp1Cre.Efnb2*^*f/f*^ bone showed a lower amide I, but higher amide II peak, suggesting greater compaction of collagen in *Dmp1Cre.Efnb2*^*f/f*^ mice.

**Figure 4:**
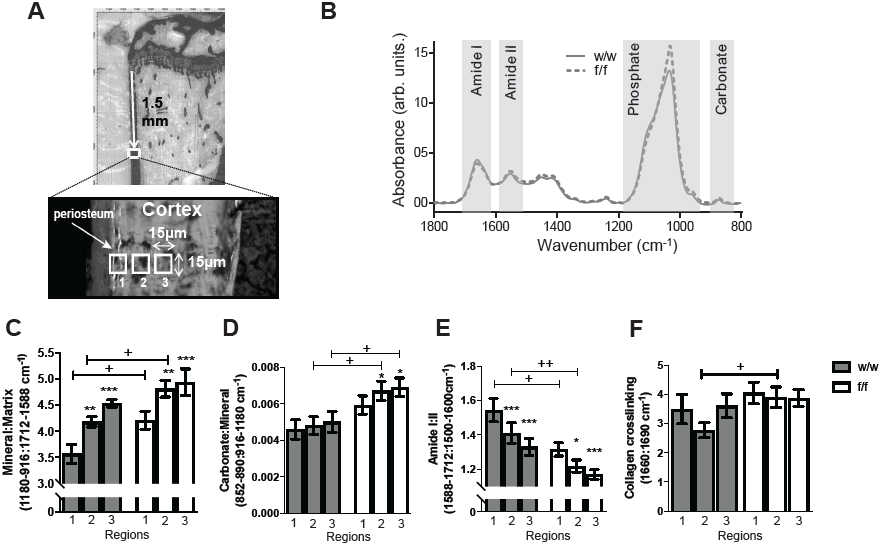
Elevated mineral accrual, carbonate deposition, and reduced amide 1:11 ratio in 12-week old female *Dmp1Cre.Efnb2*^*f/f*^ cortical bone. A: Regions of periosteal bone used for sFTIRM analysis. Note calcein label on the newly mineralized periosteum; regions 1-3 (15μm^2^ in size) denote bone areas of increasing maturity with increasing distance from the periosteal edge. B: Averaged sFTIRM spectra from region 2 of all *Dmp1 Cre.Efnb2*^*w*^*/*^*w*^ (w/w) and *Dmp1Cre.Efnb2i/i* (f/f) tibiae; boxes show approximate regions of the Amide I, Amide II, Phosphate and Carbonate peaks. C-E: sFTIRM-derived mineral ¡matrix (C), carbonate:mineral (D), amide 1:11 (E) and collagen-crosslinking (F) ratios in regions 1-3 of w/w and f/f tibiae. Data are represented as mean ± SEM, n = 13/group. *p<0.05, **p<0.01, ***p<0.001 vs region 1 of same genotype (bone maturation effect), ^+^p<0.05, ^++^p<0.01 vs w/w in the same region (genotype effect).

When quantified, control bones showed the changes associated with bone matrix maturation previously reported on murine periosteum: with increasing depth from the bone edge, mineral:matrix ratio increased while amide I:II ratio decreases ^17^ (Fig. 4C-E). The process of mineral accrual was accelerated in *Dmp1Cre.Efnb2*^*f/f*^ bone (Fig. 4C,D): in the two most immature regions of bone, mineral:matrix ratio was significantly greater in *Dmp1Cre.Efnb2*^*f/f*^ bone compared to control (Fig. 5C) and amide I:II ratio was significantly lower (Fig. 5E). In addition, although carbonate:mineral ratio did not increase significantly with maturation in the control mice, there carbonate:mineral ratio increased with increasing matrix maturity in the *Dmp1Cre.Efnb2*^*f/f*^ bone and reached a higher level of carbonate:mineral ratio than controls in the two more mature regions (Fig. 5E). The increase in mineral matrix ratio was not due to a lower bone collagen content; hydroxyproline levels were significantly higher in whole bone samples from *Dmp1Cre.Efnb2*^*f/f*^ bone compared to control (Table 3). Collagen crosslinking did not change with bone maturity, but was significantly greater in the intermediate region of *Dmp1Cre.Efnb2*^*f/f*^ bones (Fig. 4F) indicating that crosslinks mature in the same region in which mineral and carbonate accumulate. The *Dmp1Cre.Efnb2*^*f/f*^ brittle bone phenotype is therefore associated with more rapid bone matrix maturation, including rapid accumulation of mineral and carbonate substitution and more rapid collagen compaction within the matrix.

**Figure 5:**
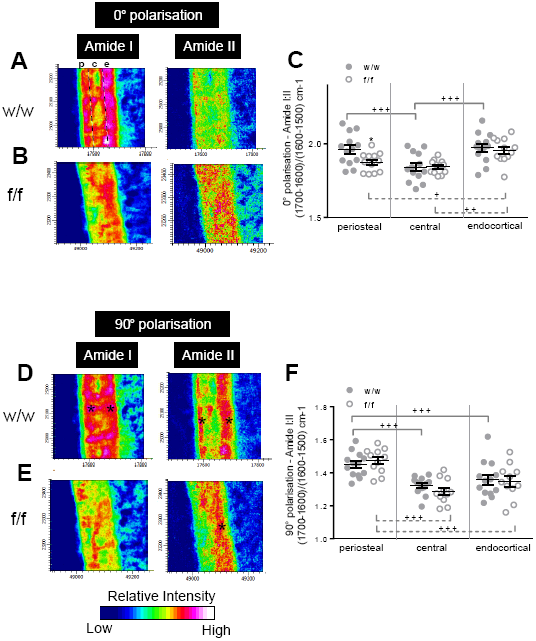
Polarised FTIR imaging confirms altered collagen distribution in 12-week old female *Dmp1Cre.Efnb2*^*f/f*^ mice. Representative FTIR images showing relative intensity of Amide I and Amide II peaks in the cortical midshaft from *Dmp1 Cre.Efnb2*^*w*^*/*^*w*^ (w/w) and *Dmp1Cre.Efnb2i/i* (f/f) mice under the (A,B) 0° and (D,E) 90° polarising filters; quantification of the amide 1:11 ratio under the (C) 0° and (F) 90° polarising filter in periosteal, intracortical and endocortical regions; regions shown in the top left panel of A; p = periosteal region, c = central region, e = endocortical region. Data are represented as mean ± SEM, n = 12-13/group. *p<0.05 vs *Dmp1Cre.Efnb2*^*w*^*/*^*w*^ at the same region (genotype effect), +p<0.05, ++p<0.01, +++p<0.001 vs region indicated by the bracket, within the same genotype (region effect).

Since the amide I:II ratio has not yet been validated in bone as an indicator of collagen orientation, we used pFTIRI in the same region to visualise and quantitate the spatial variation of collagen fibres (Fig. 5A,B,D,E). The 0° and 90° polarising filters were used to preferentially enhance the signal from molecular bonds oriented parallel and perpendicular, respectively, to the bone tissue section surface ^18^. We noted three “zones” within the cortical bone in control tibiae, where the pFTIRI 0°polarising filter indicated higher amide I:II ratio on both periosteal and endocortical regions compared to central cortical bone (Fig. 5A,C). *Dmp1Cre.Efnb2*^*f/f*^ bones showed a greater amide I:II ratio in the endocortical region compared to periosteal and central cortical bone (Fig. 5B,C). When quantified, the pFTIRI 0° polarising filter result validated the sFTIRM observations, and confirmed a significantly lower amide I:II ratio in the periosteal region in *Dmp1Cre.Efnb2*^*f/f*^ bone compared to control (Fig. 5C). In *Dmp1Cre.Efnb2*^*f/f*^ mice, the difference in amide I:II ratio between periosteal and central cortical regions was no longer detected, with both showing significantly lower amide I:II ratio than the endocortical region (Fig. 5B,C). When quantiated under the 90° polarising filter, the amide I:II ratio was significantly lower in control bone at intracortical and endocortical regions due to greater amide II signal (Fig. 5D, F). A similar observation was made in *Dmp1Cre.Efnb2*^*f/f*^ mice with the 90° polarising filter (Fig. 5E,F) and no difference was observed in fibres aligned under this filter between genotypes (Fig. 5F). This analysis validated the sFTIRM observations, and confirmed a significantly lower amide I:II ratio in *Dmp1Cre.Efnb2*^*f/f*^ bone compared to control.

### EphrinB2 deficiency in osteocytes leads to dysregulation of autophagy-related genes and increased autophagy

To identify mechanisms by which ephrinB2-deficiency alters mineral and matrix composition leading to fragile bones, RNA sequencing was performed on marrow-flushed femora from 12-week old *Dmp1Cre.Efnb2*^*f/f*^ and control mice. This revealed 782 up-regulated genes and 1024 down-regulated genes (FDR < 0.05). In the full list of differentially expressed genes, known osteocyte-specific and mineralization genes were not present (e.g. *Dmp1, Mepe, Sost, Phospho1, Enpp1, Enpp2, Fgf23).* Fig. 6A,B shows the top 30 differentially expressed genes; none have previously been associated with osteocyte function, bone mineralisation, or ephrinB2 function. We noted that a third of the top genes are associated with the process of autophagy. Six were upregulated in the ephrinB2-deficient bone: *Fam134b* ^19^, *Fbxo32* ^20,21^, *Lama2* ^22^, *Bnip3* ^23,24^, *Trim63* ^25^ and *Peg3* ^26^. The other four were downregulated: *Eps8l1*^*27*^, *Klf1*^*28*^ and *Tspo2*^*29*^ and *Unc5a* ^30^.

**Figure 6.**
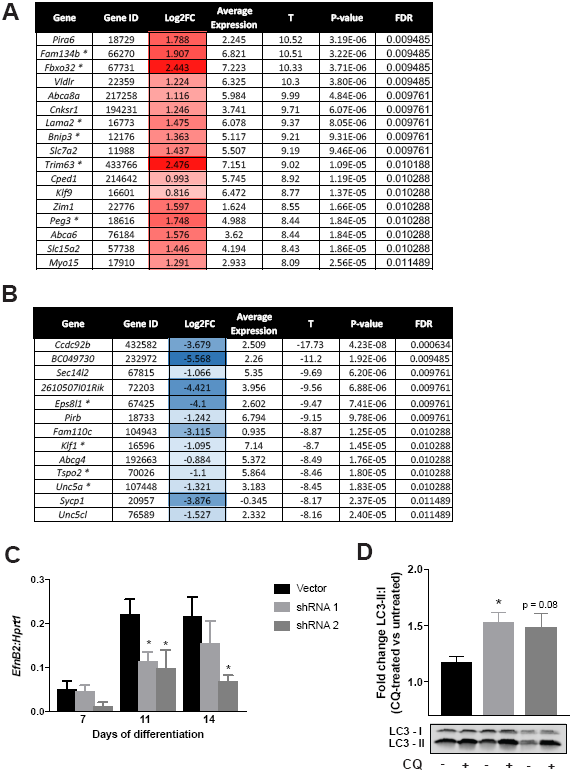
RNA sequencing from marrow-flushed femora of 12-week old female*Dmp1Cre.Efnb2*^*f/f*^ mice show upregulation of autophagy-related genes, compared to controls. Panels (A) and (B) show the top 30 most differentially expressed genes in *Dmp1Cre.Efnb2*^*f/f*^ mice compared to controls, divided into up-regulated (A) and down-regulated genes (B). Genes associated with autophagy are indicated by *. Columns give gene symbol, Entrez Gene ID, log2-fold-change, average log2-expression, moderated t-statistic, P-value and false discovery rate (FDR). C: *Efnb2* stable knockdown in Ocy454 cells differentiated for 7, 11 and 14 days, 6 replicates. D: Fold change of LC3-II:I ratio in *Efnb2* shRNA knockdown cells treated with chloroquine (CQ) for 4 hours compared to basal levels of each construct. Data are represented as mean ± SEM, 3 replicates, *p<0.05 vs. vector control.

To identify whether suppression of ephrinB2 in osteocytes leads to increased autophagy in an independent cell-specific model, we generated *Efnb2* deficient Ocy454 osteocytes and assessed autophagy levels. Stable *Efnb2* knockdown was confirmed in mature Ocy454 cells at days 11 and 14 (Fig. 6C). The rate of autophagy was measured by assessing LC3-II:I ratio after treatment with the autophagic flux inhibitor chloroquine (CQ) to allow measurement of autophagosome formation^31^. CQ treatment caused a significant elevation in LC3-II:I ratio in both vector and *Efnb2* knockdown cells, as anticipated. This effect was amplified in *Efnb2* shRNA knockdown cells, shown by the increased fold change of CQ-treated *Efnb2* knockdown cells compared to untreated cells. This indicated that ephrinB2-deficient osteocytes have greater susceptibility to autophagy than vector control (Fig. 6D).

## Discussion

This work demonstrates that bone flexibility is maintained and mineral accrual limited by ephrinB2 signalling in osteocytes. In the absence of ephrinB2, autophagic processes are increased, and although the initiation of osteoid mineralization occurs at a normal rate, as soon as the process commences, mineral deposition and maturation is accelerated, resulting in a brittle bone phenotype. This indicates that the control mechanisms for primary and secondary mineralization are different, and that osteocytes within the bone matrix control the progress of secondary mineralization. This has major implications for understanding how osteocytes contribute to bone strength.

Bone strength is controlled by both bone mass, and the composition of the bone matrix itself. Bone mass is determined by the balance between the activities of bone-forming osteoblasts and bone-resorbing osteoclasts. Bone compositional strength is determined by the content and orientation of collagen, as well as the content and nature of mineral crystals within that collagen network. Previously it has been noted that major defects in bone compositional strength can result from defective collagen deposition (as in osteogenesis imperfecta) or delayed initiation of mineralisation (as in osteomalacia or rickets). In this study, we report a third possible cause of bone fragility: accelerated maturation of the mineralized bone matrix (Fig. 7). In brittle *Dmp1Cre.Efnb2*^*f/f*^ bones, mineral matrix ratio in the first region of mineral deposition was higher than controls. Since there was no difference in the rate at which osteoid mineralization was initiated (MAR), this suggested that mineralization commenced rapidly as soon as it was initiated: essentially, mineral is “dumped” into the matrix. The normal gradual increase in mineral:matrix ratio, previously described in murine, rabbit, rat, baboon and human cortical bone ^17,32–36^, is accelerated in the absence of ephrinB2.

**Figure 7.**
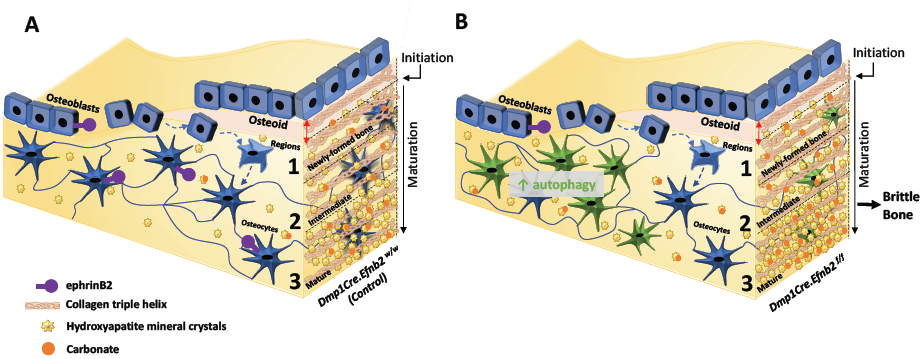
Model of how osteocytic ephrinB2 regulates bone matrix composition. (A) In control bone (*Dmp1Cre.Efnb2*^w^/^w^) both osteoblasts and osteocytes express ephrinB2. Osteoblasts reside on the bone surface, and pass through a transition (dashed arrows) to become mature, matrix embedded osteocytes. The process of bone matrix maturation is shown on the right of each 43panel, with the three measurement regions (1, 2, 3). Osteoblasts deposit collagen-containing osteoid (triple helical collagen fibres). In the first region of newly-formed bone, mineral deposition is initiated. Mineral crystals (yellow stars) and carbonte (orange circles) continue to accumulate, and collagen fibres become more compact as the matrix matures. (B) *Dmp1Cre.Efnb2*^*f/f*^ mice express ephrinB2 in osteoblasts, but not osteocytes. The transition of osteoblasts to osteocytes is increased resulting in increased osteocyte density, and osteocytes have increased levels of autophagic flux (green cells). Osteoid deposition occurs normally, and the initiation of mineralization commences at the same rate (red arrow), but as soon as it starts, mineral deposition occurs at a greater level in newly-formed bone (region 1) and in more mature bone regions (2, and 3) to reach a level where there is more mineral, more carbonate substitution and more compaction of collagen fibres compared to controls. Ultimately this leads to more brittle bone.

This was not the only aspect of mineral accumulation that was modified. Bone mineral crystals are a modified hydroxyapatite comprised of calcium, phosphate and hydroxyl ions which can be replaced by fluoride, chloride or carbonate. As hydroxyapatite matures in bone, carbonate:mineral ratio increases due to carbonate substitution for phosphate or hydroxyl ions ^17,37–40^ *Dmp1Cre.Efnb2*^*f/f*^ bones also showed a more rapid increase in carbonate incorporation within the bone matrix. Both mineral accrual and carbonate substitution within the mineral occur at an accelerated rate in the absence of ephrinB2.

A third aspect of bone matrix maturation was accelerated in *Dmp1Cre.Efnb2*^*f/f*^ bones: the reduction in the amide I:II ratio. Amide I:II ratio represents peptide bond vibrations within the collagen molecule ^41^. Amide I (C=O stretch) and amide II (C-N, N-H bend) molecular vibrations exhibit dichroism perpendicular (z-axis direction) and parallel (x-axis) to the collagen molecular triple helix axis, respectively ^42^. We have previously reported that amide I:II ratio declines with increasing mineral accumulation in control mice, likely as a result of more compaction (or steric hindrance) of the collagen molecule in the perpendicular direction as mineral accumulates ^17^. The early lowering of the amide I:II ratio in *Dmp1Cre.Efnb2*^*f/f*^ bone indicates a greater rate of collagen compaction as mineral accumulates in the matrix. This process is exaggerated in *Dmp1Cre.Efnb2*^*f/f*^ mice. Use of the 0° polarising filter suggests that increase in collagen compaction is specific to those collagen fibres aligned longitudinally to the bone section. This suggests that these fibres may be more important for absorbing mechanical forces along the length of the tibiae and their altered organization may also contribute to the bone brittleness in ephrinB2-deficient mice.

Another aspect of bone matrix maturation that was accelerated was the incorporation of osteocytes into the bone matrix: *Dmp1Cre.Efnb2*^*f/f*^ mice exhibited a greater osteocyte density. As osteoid is deposited, a subset of osteoblasts become incorporated into the bone matrix: these are the cells that differentiate into osteocytes. Our data suggests that these pre-osteocytes are incorporated into the bone matrix at a faster rate in *Dmp1Cre.Efnb2f/f* bone. *Dmp1Cre* targets late osteoblasts at the stage where they become embedded within the newly formed osteoid ^43,44^; the greater density of osteocytes incorporated may promote mineral accumulation and carbonate incorporation in *Dmp1Cre.Efnb2*^*f/f*^ bones.

Though this is the first report of a low amide I:II ratio being associated with bone fragility, the cause of the brittle phenotype is likely to be the combination of reduced amide I:II, and increased mineral:matrix and carbonate:mineral. The association of high carbonate with increased bone fragility is consistent with earlier work describing a higher carbonate:mineral ratio in bone specimens from women with postmenopausal osteoporosis ^45^ or with greater fracture susceptibility ^46^. High mineral:matrix ratio has also been reported in other examples of bone brittleness, such as patients with atypical femoral fracture ^47^, and murine models of osteogenesis imperfecta^48,49^. In the latter cases it is accompanied by defects in collagen or cartilage content and defects in osteoblast function, but the present model exhibited no change in osteoblast function.

A comparison of this mouse model with our earlier model lacking ephrinB2 throughout the osteoblast lineage provides new information about the specific stages of osteoblast/osteocyte differentiation and their roles in the process of bone mineralization. Our earlier model lacking ephrinB2 throughout the osteoblast lineage (*OsxCre.Efnb2^f/f^)* did not exhibit a brittle bone phenotype, but had bones that were more elastic due to delayed initiation of primary mineralization ^14^ This indicated that the stage of osteoblast differentiation that initiates primary mineralization is beyond the ephrinB2:EphB4 osteoblast differentiation checkpoint. In *Dmp1Cre.Efnb2*^*f/f*^ mice, osteoblasts would survive past the checkpoint at which anti-apoptotic action of ephrinB2 is required; this has allowed us to define an action for EphrinB2 later in osteoblast differentiation. Normal initiation of mineralization in *Dmp1Cre.Efnb2*^*f/f*^ bone indicates that the stage of osteoblast differentiation that controls this process is not only after the ephrinB2:EphB4 checkpoint, but before the stage of *Dmp1Cre* expression. The stage of osteoblast/osteocyte differentiation after *Dmp1* is expressed does not control initiation of mineralisation but controls the rate at which mineral accumulates (Fig. 7). Since PTH and PTHrP promote ephrinB2 expression in both osteoblasts^12^ and osteocytes, they may promote both the initiation of osteoid mineralization and restrain the rate of mineral accrual. While there is not evidence for altered mineral accrual with pharmacological administration of PTH, the reduced material strength of mice with PTHrP deletion targeted to osteocytes provides support for a role of PTHrP in regulating mineral accrual via ephrinB2-dependent actions in the osteocyte ^50^.

The processes that control the rate of mineral accumulation in bone are poorly defined and our data suggests that autophagic processes in the osteocyte that are inhibited by ephrinB2 limit this process. Unbiased RNA sequencing detected no changes in mRNA levels of genes known to regulate the initiation of mineralization *(Dmp1* and *Mepe).* Instead, it identified dysregulation of a range of autophagy-associated genes and we confirmed that ephrinB2 inhibits autophagy *in vitro.* Autophagy is a lysosome-based pathway that contributes to diverse cellular functions, such as adaptation to starvation, quality control of intracellular processes, protein secretion, and elimination of intracellular microbes ^51‐53^. While explorations of the role of autophagy in osteocytes is in its infancy, previous work has demonstrated that autophagy increases during osteoblast differentiation ^54^ and that mice with osteoblast-lineage or osteocytic deletion of either *Atg5* or *Atg7* have reduced autophagy and reduced osteoblast numbers ^54‐56^. In our *Dmp1Cre.Efnb2*^*f/f*^ model osteoblast numbers were not altered, and we observed no changes in *Atg5* or *Atg7,* suggesting a subset of autophagic processes that control mineral maturation without controlling osteoblast differentiation are modified in the absence of ephrinB2.

We suggest two ways by which increased autophagic flux may cause the brittle bone phenotype: (1) controlling osteocyte differentiation and adaptation to the mineralized environment, and (2) contributing to mineral secretion. After osteocytes are embedded in osteoid, they develop dendritic extensions and adjust to a more hypoxic environment ^57^. Autophagic processes may contribute to this transition by allowing the osteocyte to break down complex molecules to use for energy ^58^, promoting faster organelle recycling, and preservation of nutrients. Mineralization involves the release of cell-derived extracellular membrane-enclosed particles of poorly crystalline mineral, termed matrix vesicles ^59,60^ and subsequent nucleation (ordered crystal formation) of that amorphous mineral by heterogeneous nucleation driven by contact with collagen, and by secretion of competent apatite nucleators ^61‐63^. Nollet *et. al* suggested that autophagic vacuoles could serve as vehicles for osteoblasts to secrete apatite crystals in the extracellular space via exocytotic processes ^54^; we propose that osteocytes may also release mineral through this process. Direct release of mineral by osteocytes has been noted both during remineralization of osteocyte lacunae after lactation ^64^ and in egg-laying hens ^65^. Our data suggest that the secretion of both vesicles and nucleators by osteocytes promotes mineral accrual during physiological bone formation. We suggest that ephrinB2-deficient osteocytes use autophagic machinery to release more matrix vesicles and apatite nucleators, thereby driving an increase in bone mineral, resulting in brittle bone matrix.

These findings have implications for other organ systems. EphrinB2 in odontoblasts, which are closely related cells to osteoblasts and osteocytes ^66,67^ may also control mineralisation in teeth. Pathological mineralisation, such as that in heterotopic ossification after trauma ^68^, vascular or renal calcification ^69,70^ may also involve autophagy and be limited by ephrinB2, since it is also expressed in those tissues.

In conclusion, ephrinB2 is required to restrain autophagy in the osteocyte, and to prevent the formation of brittle bone. Osteocytic ephrinB2 limits the accumulation of mineral and carbonate substitution within the hydroxyapatite matrix and restrains collagen fibre compaction. This indicates a mechanism by which bone strength is regulated independently of bone size, and independently of osteoblast and osteoclast activities.

## Experimental Procedures

### Cell culture

Ocy454 cells were cultured as previously described, in αMEM supplemented with 10% FBS and 1% Penicillin-Streptomycin-Amphotericin B and Glutamax ^71^. Cells were maintained in permissive conditions (33°C) and differentiated to osteocytes at 37 °C ^71^. For 2D cell cultures, Ocy454 cells were plated at 2.5 x 10^5^ cells per well in six-well plates. Three-dimensional (3D) culture was performed using Reinnervate^®^ Alvetex scaffold 6-well inserts (Pittsburgh, PA, USA) made of highly porous, cross-linked polystyrene discs with 200 μm thickness and 22mm diameter. The Alvetex inserts were prepared for seeding by a 70% ethanol wash, followed by two washes with complete culture media. Cells were seeded at 1.6 x 10^6^ cells per insert. Cells were grown at the permissive temperature (33°C) for 3 days prior to transferring to (37°C) for differentiation.

To study how *Efnb2* mRNA is regulated by PTH and PTHrP, Ocy454 cells at day 14 of differentiation were serum starved overnight in aMEM supplemented with 1% FBS and 1% PSA and Glutamax. Cells were then treated with Human PTH(1-34) (10 nM) or Human PTHrP(1-141) (10 nM) for 6 hours. After 6 hours, cells were washed with phosphate buffered saline (PBS) and RNA samples were collected as described below.

### Pthlh knockdown

shRNA with sequence 5’CCG-GCC-AAT-TAT-TCC-TGT-CAC-TGT-TCT-CGA-GAA-CAG-TGA-CAG-GAA-TAA-TTG-GTT-TTT-TG-3’was used to knock down *Pthlh* in Ocy454 cells as previously reported ^9^. Undifferentiated Ocy454 cells were infected with virus and selected with puromycin (5 ug/ml), then cultured at permissive temperature (33°C) before transfer to 37°C for differentiation. Cells were assessed by quantitative real-time PCR (qRT-PCR) at day 0 (undifferentiated), 7 and 14 (differentiated)^9^.

### qRT-PCR analysis

RNA was extracted by RNA extraction kits with on-column DNase digestion (Qiagen, Limburg, Netherlands; Bioline, London, UK), or TriSure reagent (Bioline, London, UK). Extracted RNA was DNase treated with Ambion TURBO DNA-free kit (Life Technologies) and quantified on a NanoDrop ND1000 Spectrophotometer (Thermo Scientific, Wilmington, DE, USA). cDNA was synthesised from total RNA with AffinityScript cDNA synthesis kits (Agilent Technologies, Santa Clara, CA, USA). Gene expression for PTH and PTHrP treated Ocy454 cells were quantified on a Stratagene Mx3000P QPCR system (Agilent) with SYBR Select Master Mix (Applied Biosystems) with primers specific to *Efnb2;* forward 5’-GTGCCAGACAAGAGCCATGAA-3’ and reverse 5’-GGTGCTAGAACCTGGATTTGG-3’^12^. Gene expression *for Pthlh* knockdown cells was analysed using the Multiplex SensiMix II Probe kits (Bioline, London, UK) with primers specific to *Efnb2,* as previously described ^13^. Gene expression levels between samples was normalized to hypoxanthine phosphoribosyltransferase 1 (*Hprt1)* expression. Relative expression was quantified using the comparative CT method (2^-(Gene Ct-Normalizer Ct)^).

### Mice

Dmp1Cre mice (Tg(Dmp1-Cre)^1Jqfe^) (containing the DMP1 10-kb promoter region) were obtained from Lynda Bonewald (University of Kansas, Kansas City, KS, USA) ^43^ and ephrinB2-floxed (Efnb2^tm1And^) mice were obtained from David J. Anderson (Howard Hughes Medical Institute, California Institute of Technology, Pasadena, CA, USA) ^72^; all mice were backcrossed onto C57BL/6 background. Mice hemizygous for Dmp1Cre were crossed with Efnb2^f^/^f^ mice to generate Dmp1Cre.Efnb2^f^/^w^ breeders, which were used to generate Dmp1Cre.Efnb2^f^/^f^ mice and Dmp1Cre.Efnb2^w^/^w^ littermates or cousins, which were used as controls for all experiments. To generate GFP+ osteocytes, these mice were crossed with Dmp1-GFP (Tg(Dmp 1-Topaz)^1Ikal^ mice obtained from Dr Ivo Kalajzic, University of Connecticut Health Science Center, via the colony of Dr Hong Zhou, ANZAC Research Institute, Sydney ^73^. For all tissue collections, mice were fasted for 12 hours prior to anaesthesia with ketamine and blood was collected via cardiac puncture, as previously described ^74^ Both male and female mice were analysed. All animal procedures were conducted with approval from the St. Vincent’s Health Melbourne Animal Ethics Committee.

### Confirmation of Efnb2 mRNA targeting in osteocytes

To confirm specific targeting of *Efnb2* mRNA in osteocytes, *Dmp1Cre.Efnb2*^*f/f*^ mice were crossed with *Dmp1-GFP* mice to allow purification of osteocytes by fluorescence-activated cell sorting (FACS), according to our previously published methods ^9,75^. Osteocytes from 6-week old *Dmp1Cre.Dmp1-GFP-Tg.Efnb2*^*w*^*/*^*w*^ and *Dmp1Cre.Dmp1-GFP-Tg.Efnb2*^*f/f*^ mice were isolated from marrow-flushed long bones by seven sequential 15-minute digestions in 2 mg/ml dispase (Gibco, Grand Island, NY, USA) and 1 mg/ml collagenase type II (Worthington, Lakewood, NJ, USA). Fractions 2-7 were collected, pooled, and resuspended in alpha modified Eagle’s medium (α-MEM; Gibco, Grand Island, NY, USA) containing 10% FBS and centrifuged. Pellets were resuspended in FACS buffer before cell sorting. Prior to sorting, dead cells and debris were removed based on side scatter (SSC) area and forward scatter (FSC) area, and doublets were excluded based both on SSC width (W) vs SSC height (H) and on FSC-W vs FSC-H. Cells were sorted with excitation 488 nm and 530/30 or 530/40-emission filter for GFP on a BD FACS Influx cell sorter (BD Biosciences, Scoresby, Australia). RNA was extracted using Isolate II Micro RNA kit (Bioline, London, UK). cDNA was prepared using a Superscript III kit (Thermo Fisher, Scoresby, Australia). *Efnb2* primers directed to the targeted region were: forward 5’-AGAACTGGGAGCGGCTTG-3’, reverse 5’-TGGCCAACAGTTTTAGAGTCC-3’^14^. Gene expression levels between samples were normalised to Beta-2 microglobulin (*β2m*): forward 5’-TTC ACC CCC ACT GAG ACT GAT-3’, reverse 5’-GTC TTG GGC TCG GCC ATA-3’ and hydroxymethylbilane synthase (*Hmbs*): forward 5’-TCATGTCCGGTAACGGCG-3’, reverse 5’-CACTCGAATCACCCTCATCTTTG-3’-expression. Relative expression was quantified using the comparative threshold cycle (Ct) method (2^-(Gene Ct-Normalizer Ct)^).

### 3-point bending and Reference Point Indentation (RPI)

Structural and material properties were analysed in femora from male and female 12-week-old mice by 3-point bending, as described previously ^74^ Cortical dimensions including anteroposterior (AP) and mediolateral (ML) widths were measured at the cortical midshaft by microCT. Moment of inertia was calculated based on AP, ML and cortical thickness, as previously described ^9^. Load was applied in the anterior-posterior direction of the femoral mid-shaft between two supports that were 6.0mm apart. Load-displacement curves were recorded at a crosshead speed of 1.0 mm/s using an Instron 5564A dual column material testing system, and Bluehill 2 software (Instron, Norwood, MA, USA). Ultimate force (N), ultimate deformation (mm), post-yield deformation (mm) and energy absorbed to failure (mJ) were measured from the load-displacement curves. Combining the geometric calculations and the biomechanical test results, the material properties of each bone were calculated as previously described ^74^ to obtain ultimate stress (MPa), ultimate strain (%) and toughness (J/mm^3^).

Local bone material properties were examined by reference point indentation (RPI) using a BP2 probe assembly apparatus (Biodent Hfc; Active Life Scientific Inc., Santa Barbara, CA, USA), as previously described ^74^ Briefly, measurements were taken at the femoral mid-shaft, 6 mm from the base of the femoral condyles for consistent probe positioning between samples. A maximum indentation force of 2N was achieved by manually applying a 300g reference force to femora. 5 measurements were taken per sample using a 2N, 10-cycle indentation protocol at 2N force. Pre-and post-experiment measurements were taken on a polymerized methyl methacrylate block to ensure that probe assembly was not affected during testing. The indentation distance increase (IDI) was measured as the indentation distance in the last cycle relative to the first cycle ^76^.

### Histomorphometry, back scatter electron microscopy (BSEM) and Micro-Computed Tomography (microCT)

Tibiae from 12-week-old female mice were fixed in 4% paraformaldehyde, embedded in methyl methacrylate (MMA), sectioned longitudinally and stained as described previously ^13^. Calcein was administered by intraperitoneal (IP) injection 7 and 2 days before tissue collection. Periosteal parameters were measured on the medial side of the tibial mid-shaft (1500 μm from the base of the growth plate), as previously described ^77^. Trabecular parameters were measured in the secondary spongiosa, commencing 370 μm below the growth plate, in a 1110 μm^2^ region in the proximal tibia, as previously described ^77^ (Osteomeasure; Osteometrics, Atlanta, GA, USA).

Polarized light microscopy and BSEM were performed on 100 μm thick transverse sections from the femoral mid-shaft generated using an Isomet Saw (Buehler, Lake Bluff, IL, USA); measurements included the entire bone interface. Irregular fibre orientation was measured as woven bone and well-aligned fibre orientation was measured as lamellar bone, as previously described ^74^.

BSEM was used to acquire gray-level images of mineralized cortical bone to identify osteocyte lacunae. Osteocyte lacunae were imaged on both anterior and posterior sides of the femur at 500x magnification (approximately 600 μm in length) using an FEI Quanta FEG 200 solid state backscattered scanning electron microscope. Imaging was performed at low vacuum using water at a 9.8mm working distance. Lacunae were quantified using MetaMorph (v7.8.3.0; Molecular Devices; Sunnyvale, CA, USA) by establishing an inclusive threshold for dark objects to distinguish bone from background. Integrated Morphometry Analysis (IMA) was applied with a 2.43 - 24.31 μm^2^ filter for lacunae area, to exclude cracks and blood vessels. The thresholded bone area was combined with total lacunar area to calculate the total bone area. Parameters measured included osteocyte lacunar size, size of the largest 20% of osteocyte lacunae and osteocyte lacunar density.

MicroCT was performed on femora using the SkyScan 1076 system (Bruker-microCT, Kontich, Belgium). Images were acquired using the following settings: 9μm voxel resolution, 0.5mm aluminium filter, 50 kV voltage, and 100 μA current, 2600 ms exposure time, rotation 0.5 degrees, frame averaging =1. Images were reconstructed and analysed using NRecon (version 1.6.9.8), Dataviewer (version 1.4.4) and CT Analyser (version 1.11.8.0). Femoral trabecular analysis region of interest (ROI) was determined by identifying the distal end of the femur and calculating 10% of the total femur length towards the mid-shaft, where an ROI of 15% of the total femur length was analysed. Bone structure was measured using adaptive thresholding (mean of min and max values) in CT Analyser. Cortical analyses were performed in a region of 15% of the femoral length commencing from 30% proximal to the distal end of the femur and extending toward the femoral mid-shaft. The lower thresholds used for trabecular and cortical analysis were equivalent to 0.20g/mm^3^ and 0.642g/mm^3^ calcium hydroxyapatite (CaHA), respectively. Cortical tissue mineral density (Ct.TMD) was analysed in the same cortical region of interest, as described above. TMD calibration was performed using two phantom rods with concentrations of CaHA of 0.25 and 0.75g/cm^−3^, scanned under the same conditions and settings as the samples.

### Synchrotron-based Fourier Transform Infrared Microspectroscopy (sFTIRM) and polarized light FTIR Imaging (pFTIRI)

sFTIRM was used to examine bone composition at the cortical diaphysis in 3μm longitudinal sections of methyl methacrylate (MMA)-embedded tibiae. Sections were imaged using a Bruker Hyperion 2000 IR microscope coupled to a V80v FTIR spectrometer located at the IR Microspectroscopy beamline at the Australian Synchrotron, as previously described ^*17*^. Sections were placed on 22mm diameter x 0.5mm polished barium fluoride (BaF2) windows (Crystan Limited, UK). The microscope video camera was used to image the cortical diaphysis (1.5 mm from the base of the growth plate), at the same location used for histomorphometric measurements of periosteal mineral apposition (Fig 5A). This location is ideal for assessing mineral apposition on a formation surface, without any prior remodelling because mouse bone lacks haversian systems ^78^. sFTIRM mapping was performed with the synchrotron source, using a 15 x 15 μm aperture, represented by the 3 regions within the cortex. Spectra were collected from these 3 regions which progressed perpendicularly into the cortex with the first positioned at the periosteal edge (Fig 5A). Spectra were collected in the mid-IR region from 750 cm^−1^ to 3850 cm^−1^ using a narrowband mercury cadmium telluride detector, at 8 cm^−1^ spectral resolution and 128 co-added scans per pixel spectral resolution in transmission mode. A matching background spectrum was collected through clear BaF2. For each sample MMA reference spectra were collected from within the embedding material. All data acquisition was undertaken with Bruker OPUS version 6.5 and data analysis completed with OPUS version 7.2.

After acquisition, raw spectra for each region and sample were baseline corrected using a 3-point baseline at 1800, 1200 and 800 cm^−1^. Residual MMA absorbance peaks were then subtracted using the relevant MMA reference spectrum for each sample by iterative manual subtraction. A residual 1730 cm^−1^ MMA band remained after MMA subtraction which was not used for analysis. Spectroscopic parameters calculated were integrated peaks areas of the following bands: Phosphate (1180-916 cm^−1^), amide I (1588-1712 cm^−1^) and II (1600-1500 cm^−1^), carbonate (890-852 cm^−1^), Ratios were calculated as follows: mineral:matrix ratio (1180-916cm^−1^/588-1712 cm^−1^), carbonate:phosphate ratio (1180-916cm^−1^/890-852 cm^−1^) ratio and amide I:II ratio (1588-1712 cm^−1^/1600-1500 cm^−1^) ^*79*^*-*^*80*^. Collagen crosslinking was determined by spectral curve fitting of the amide I and amide II peaks using Grams/AI (Version 9.2, Thermo Scientific, USA). The second-derivative of each peak was used to estimate subpeak positions at approximately 1660cm^−1^ and 1690cm^−1^, as previously described ^17^.

pFTIRI was applied to the same 3μm tibial sections at the same cortical diaphyseal region of analysis used for sFTIRM. This time, larger regions were imaged using a 340 x 340μm aperture either side of the 1500 μm mid-point. Tissue sections were scanned using a Hyperion 3000 spectral imaging system equipped with a Vertex-70 spectrometer (Bruker, Germany), a liquid-N2 cooled focal plane array (FPA: 128×128 elements; 40×40 microns each) detector and a Globar source. The FPA detector was continuously maintained at liquid-N2 temperature by an automated refilling system (Norhof LN2-cooling system #606; Maarssen, Netherland). The microscope and spectrometer were also continuously N2-purged and an insulation box protected the sample stage from ambient air. For all FTIR image acquisitions, a 15x magnification level and condenser were used. High-resolution FTIR images were obtained for microscopic analysis of tissue sections; 200 scans and an 8 cm^−1^ spectral resolution was used for image acquisitions (140 ms FPA detector exposure per scan; spectral range = 3800-900 cm^−1^). All FTIR images had an individual pixel dimension of 2.6×2.6 μm, thus at ∼λ/2 for the mid-IR spectral interval. All infrared images were obtained in transmission mode. The images were obtained from sub-routines of the Opus 7.5 software (Bruker-Optics, France). To quantitate region specific changes in amide I:II, the diaphyseal cortex was divided into thirds to obtain images of periosteal, intracortical and endosteal regions of interest. Images were quantitated using CytoSpec 1.4.0.3 (Bruker) and average spectra were extracted from each region, baselined and MMA-subtracted (similar to above). Integrations for the amide I and II peaks were calculated as follows: amide I (1590-1730 cm^−1^) and amide II (921-1190 cm^−1^). We interpreted the amide I:II ratio as peptide bond vibrations of the collagen molecule. The amide I (C=O stretch) and amide II (C-N, N-H bend) molecular vibrations exhibit dichroism perpendicular (z-axis direction) and parallel (x-axis) to the collagen molecular triple helix axis, respectively ^18,41^. The FTIR microscope was coupled with two polarising filters to measure molecular orientation of collagen fibres relative to the plane of the tissue section. The 0° polarising filter was used to measure bonds “in plane” (parallel to the section). The 90° polarising filter was used to measure bonds “out of plane” (perpendicular to the section). This allowed quantification of collagen fibres aligned in different directions through the diaphyseal cortex.

### Hydroxyproline assay in hydrolysed bone samples

Following mechanical testing, all fragments of the broken femora were flushed of marrow, hydrolyzed and used for hydroxyproline assay to measure collagen content (31). Femoral samples were cut with scissors at the mid-diaphysis and the distal half of the femur flushed of marrow. The bone was weighed before and after dehydration in a 37°C incubator for one hour. Each sample was hydrolyzed by incubation overnight at 120°C in 1mL of 5M HCl per 20mg of bone. 50μl of each sample was used, and 6 serial dilutions of the hydroxyproline standard (1mg/mL) were used to generate a standard curve. 50μl of Chloramine T (Sigma-Aldrich) was added to each reaction and incubated at room temperature for 25 minutes. 500μl of Ehrlich’s Reagent (Dimethyl-amino-benzaldehyde (Sigma-Aldrich), n-propanol (99%, Sigma-Aldrich), perchloric acid (70%, AnalAR)) was added to each reaction and incubated at 65°C for 10 minutes (or longer if colour was not well developed). 100μl of each reaction was pipetted into a 96 well plate and absorbance measured using a POLARstar plate reader at 550nm and interpolated on the standard curve ^81^.

### RNA sequencing

RNA samples were collected from flushed femora of 12 -week old female *Dmp1Cre.Efnb2*^*f/f*^ and cousin-bred control mice. Samples were collected on two occasions: on day 1, two *Dmp1Cre.efnB2f/f* and three control samples were collected, while on day 2, one sample of each genotype was collected. Bones were snap frozn in liquid nitrogen, homogenized in QIAzol lysis reagent with a Polytron PTA 20S homogenizer at 4C prior to RNA extraction with a RNeasy Lipid Minikit (QIAgen). RNA sequencing was conducted on an Illumina HiSeq at the Australian Genome Research Facility to produce 100bp paired-end reads. Reads were mapped to the mouse genome (mm10) using Rsubread ^82^. Read counts were obtained using featureCounts and Rsubread’s inbuilt mm10 annotation ^83^. Gene annotation was obtained from the NCBI gene information file downloaded 4 October 2016. Statistical analysis used the limma software package ^84^. Genes were retained in the analysis if they achieved at least 0.65 read counts per million (cpm) in at least 3 samples. Immunoglobulin gene segments, ribosomal genes, predicted and pseudo genes, sex-linked genes (Y chromosome and Xist) and obsolete Entrez Gene IDs were filtered out. Quantile normalization was applied and read counts were transformed to log2-counts-per-million. Linear models were used to test for expression differences between deficient vs. control samples. The day of sample collection was included in the linear model as a blocking factor. Empirical sample quality weights were estimated ^85^. Differential expression between the genotypes was assessed using empirical Bayes moderated t-statistics allowing for an abundance trend in the standard errors and for robust estimation of the Bayesian hyperparameters ^86^. The Benjamini and Hochberg method was used to adjust the p-values so as to control the false discovery rate.

### shRNA knockdown of ephrinB2 in Ocy454 cells

Two shRNA constructs were used to knock down ephrinB2: shRNA 1 (5’-CGG-GTG-TTA-CAG-TAG-CCT-TAT-3’) and shRNA 2 (5’-CAG-ATT-GTG-TAC-ATA-GAG-CAA-T-3’), both obtained from Sigma-Aldrich (St. Louis, MO, USA). shRNA were cloned into the PLKO lentiviral vector, as previously reported ^9^ and infected by retrovirus into undifferentiated Ocy454 cells. Infected cells were selected with puromycin (5μg/mL) and cultured at permissive temperature (33°C) before transfer to 37°C for differentiation. Knockdown was validated by quantitative real-time PCR (qRT-PCR).

### Western blot analysis of chloroquine-treated Ocy454 cells

To test the level of autophagy in osteocytes lacking *Efnb2,* Ocy454 cells with *Efnb2* shRNA knockdown and vector controls were differentiated for 11 days and treated with chloroquine (40μm), a lysosomal degradation inhibitor, for 4 hours. Treated and untreated cells were then washed with phosphate-buffered saline (PBS) and lysed in 50 mM Tris, pH 7.4, 150 mM NaCl, 10% glycerol, 1 mM EDTA, 1 mM EGTA, 0.1% NaPyroP, 1% Triton-X-100, Roche protease inhibitor. Samples were electrophoresed by 6% SDS-PAGE and transferred to Immobilon FL polyvinylidine-flouride membrane (Millipore). Membranes were blotted with antibodies raised against LC3B (D11, Cell Signaling Technology, 3868), followed by incubation with anti-rabbit IgG 2° antibody fluorescently labelled with IR680 (LI-COR Biosciences). Immunoblots were visualised on an Odyssey membrane imaging system (LI-COR Biosciences) and the LC3II:I ratio was quantitated as the fold change of chloroquine-treated samples relative to untreated samples.

### Statistics

All graphs show mean ± SEM; number of samples (n) is reported in the figure legends. Statistical significance was determined by unpaired Student’s t-tests for histomorphometric, mechanical and microCT analysis, and two-way ANOVA with Fisher’s LSD test for sFTIRM-derived data (GraphPad Prism 6 (version 6.05)). p<0.05 was considered statistically significant.

## Acknowledgements

The authors thank the staff of the St. Vincent’s Health Bioresources Centre for excellent animal care and assistance, Mr Joshua Johnson and Mrs Ingrid Poulton for technical assistance with histology, Dr Roger Curtain (Bio21) for technical assistance with BSEM, Dr Paul Roschger for advice on BSEM analysis and Dr Eleftherios Paschalis for advice on collagen crosslinking analysis. This work was supported by NHMRC Grants 1042129 and 1081242 to NAS and TJM, Program Grant 1054618 to GKS, the Australian and New Zealand Bone and Mineral Society Christine and TJ Martin Travel Award to CV, and a Brockhoff Foundation Grant to CV. JSO was supported by an ARC Future Fellowship. NAS was supported by an NHMRC Senior Research Fellowship. St. Vincent’s Institute acknowledges the support of the Victorian State Government OIS program. Part of this work was untaken at the Infrared Microspectroscopy Beamline at the Australian Synchrotron, part of ANSTO.

## Author contributions

Conceptualization, C.V., T.J.M. and N.A.S.; Investigation, C.V., T.A.D., N.A., B.C.I., H.N., Y.H., M.I., G.K.S., N.A.S.; Data curation, C.V., Y.H., G.K.S.; Analysis, C.V., T.A.D., N.A., C.P., Y.H.; Writing - Original Draft, C.V., N.A.S.; Writing-Review & Editing, all authors; Visualization, C.V., N.A.S.; Supervision, N.A.S.; Funding acquisition, N.A.S.

## Declaration of Interests

The authors declare no competing interests.

